# Cementochronology using synchrotron radiation tomography to determine age at death and developmental rate in the holotype of *Homo luzonensis*

**DOI:** 10.1101/2023.02.13.528294

**Authors:** Anneke H. van Heteren, Andrew King, Felisa Berenguer, Armand Salvador Mijares, Florent Détroit

## Abstract

*Homo luzonensis*, a fossil hominin from the Philippines, is smaller than modern humans. At present, very little is known about the life history of this species. Cementochronology can answer life history questions, but usually involves destructive sampling. Here, we use synchrotron radiation to count the yearly cement lines of teeth belonging to a single individual. This approach allows us to determine that this individual was likely 31 years old at time of death and apparently had a developmental pattern comparable to chimpanzees. To our knowledge, this is the first time that cementochronology using synchrotron radiation tomography is used for life history and age-at-death estimation.

## Introduction

Islands provide amazing examples of evolution and have inspired both Charles Darwin and Alfred Russel Wallace. Nonetheless, little is known about insular evolution, often resulting in miniature or giant versions of mainland animals (1). Whether hominins are also subject to the insular dwarfing to the same degree as other taxa, is still a matter of debate (2, 3) and possible examples include the *Homo floresiensis* from Indonesia (4) and *Homo luzonensis* from the Philippines (5).

The remains of *Homo luzonensis* were found in Callao Cave on Luzon Island and were described in 2019 (6). The holotype of the species consists, amongst other bones and teeth, of a dental series from upper right P^3^ to upper right M^3^, CCH6-a to CCH6-e, (6). The teeth are characterised by their small size and mosaic of primitive and derived features (7). Two *Homo luzonensis* elements, the third metatarsal CCH1 and the upper right M3 CCH6-a, were directly dated to minimum ages of 67 thousand years (kyr) and 50 kyr respectively (5, 8). Whether these hominin remains represent a species that dwarfed under the influence of insular conditions remains to be determined (6), although a descent from *Homo erectus* seems likely based on the internal organisation of its teeth (7). This begs the questions: Did the life history parameters of this species differ from that of other hominins? And, how old was this individual when he or she died?

In medical terms, the incremental lines in tooth cementum are called “lines of Salter”(9, 10). The morphology of tooth cementum resembles lamellar bone (11). As in the case of lamellar bone (12), there are heavy debates on the nature and the depositional mechanism of the lines of Salter in tooth cementum (13) centering around analogous hypotheses. It is generally agreed upon that the lines of Salter are annual lines representing summer and winter (13). Nevertheless, the study of seasonality in human cementochronology has mostly been performed on samples from the northern hemisphere from areas with four seasons (13). It is still not understood how the colours of the lines of Salter correspond to the seasons in the southern hemisphere or in areas with monsoonal seasonality.

There is no evidence for tooth cement abrasion during decomposition, unlike tooth enamel (14), making tooth cement a promising candidate for chronological research. No taphonomical studies on geological time scales have thus far been performed, however. In the present paper, we answer the above research questions using cementochronology, or tooth cementum annulation, derived from synchrotron radiation tomography.

## Materials and Methods

In 2015, three teeth (CCH6-d: P^4^; CCH6-c: M^1^; CCH6-b: M^2^) of the holotype of *Homo luzonensis* were scanned in Synchrotron SOLEIL in France at the PSICHE beamline. Although other researchers have, in the meantime, also applied cementochronology to modern human canines using synchrotron radiation (15, 16), this is the first instance, to our knowledge, that synchrotron based cementochronology is used on 1) a series of teeth belonging to the same individual, 2) that are completely fossilized, and 3) for the purpose of estimating life history traits through dental development.

Since transverse sections are advised for human teeth (13), P^4^ and M^2^ were mapped transversally by performing a mosaic of quick acquisitions, for which the detector image was binned 4×4 to create an effective pixel size of 2.6 microns. Each projection required 0.4 s exposure time, and 300 images were acquired over 180° (roughly 2 minutes per tomogram). The reconstructed slices were used to provide a preview of the transverse section, in order to select the locations with the best-preserved cement structures (Figure 1). The buccal root of P^4^ is broken off, so the lingual root was mapped. Rather than use a mosaic mapping approach, M^1^ was scanned, also transversally, at a relatively low resolution with 2.6 micron pixel size, 1500 projections over 180° with 170 mm propagation distance and 0.15s exposure time at 40 keV (Figure 2).

**Figure 1:**
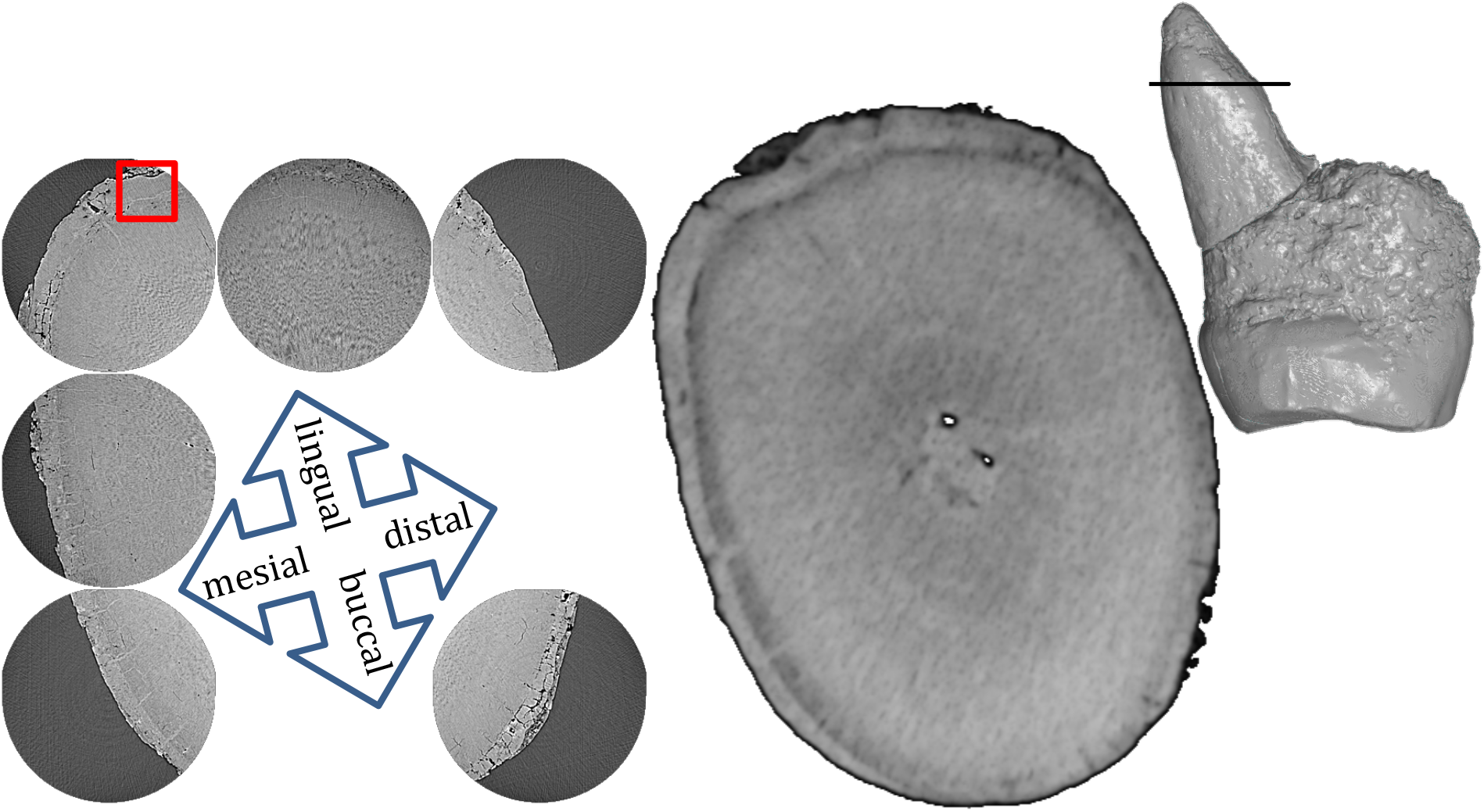
Left: Map at 6 mm from the tip of the root of P^4^(CCH6-d). The red square indicates the position of the detailed scan in Figure 3 at the lingual aspect of the lingual root. Centre: For illustrative purposes, a slice of a micro-CT scan of P^4^ (7) at the level of the map. Right: The line in the distal view of the 3D model indicates the level where the scan was made.

**Figure 2:**
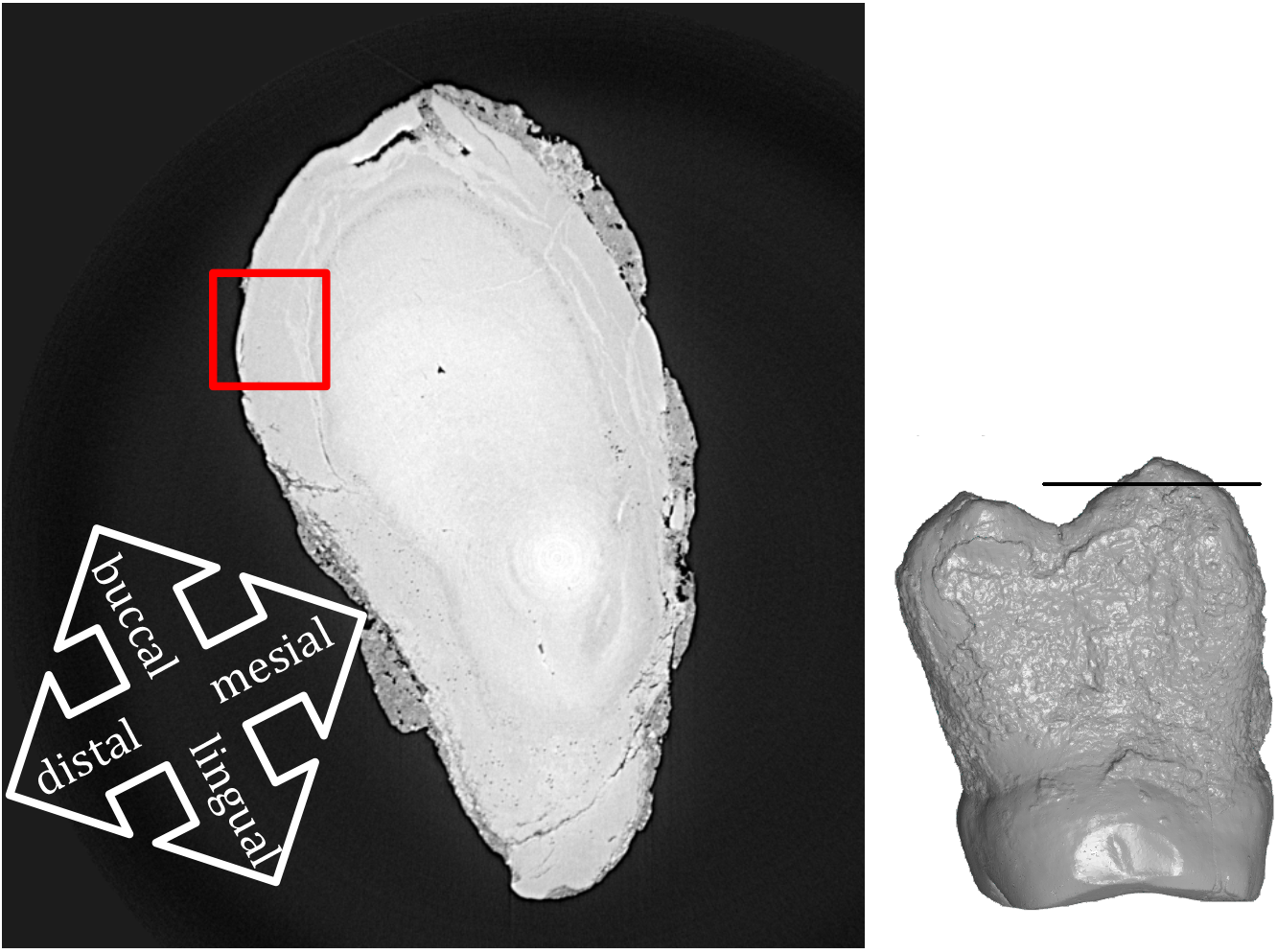
Left: Section at 0.93 mm from the buccal tip of the distal root of M^1^ (CCH6-c) on a low-resolution scan. The red square indicates the position of the detailed scan of the distobuccal aspect in Figure 4. Right: The line in the distal view of the 3D model indicates the level where the scan was made.

**Figure 3:**
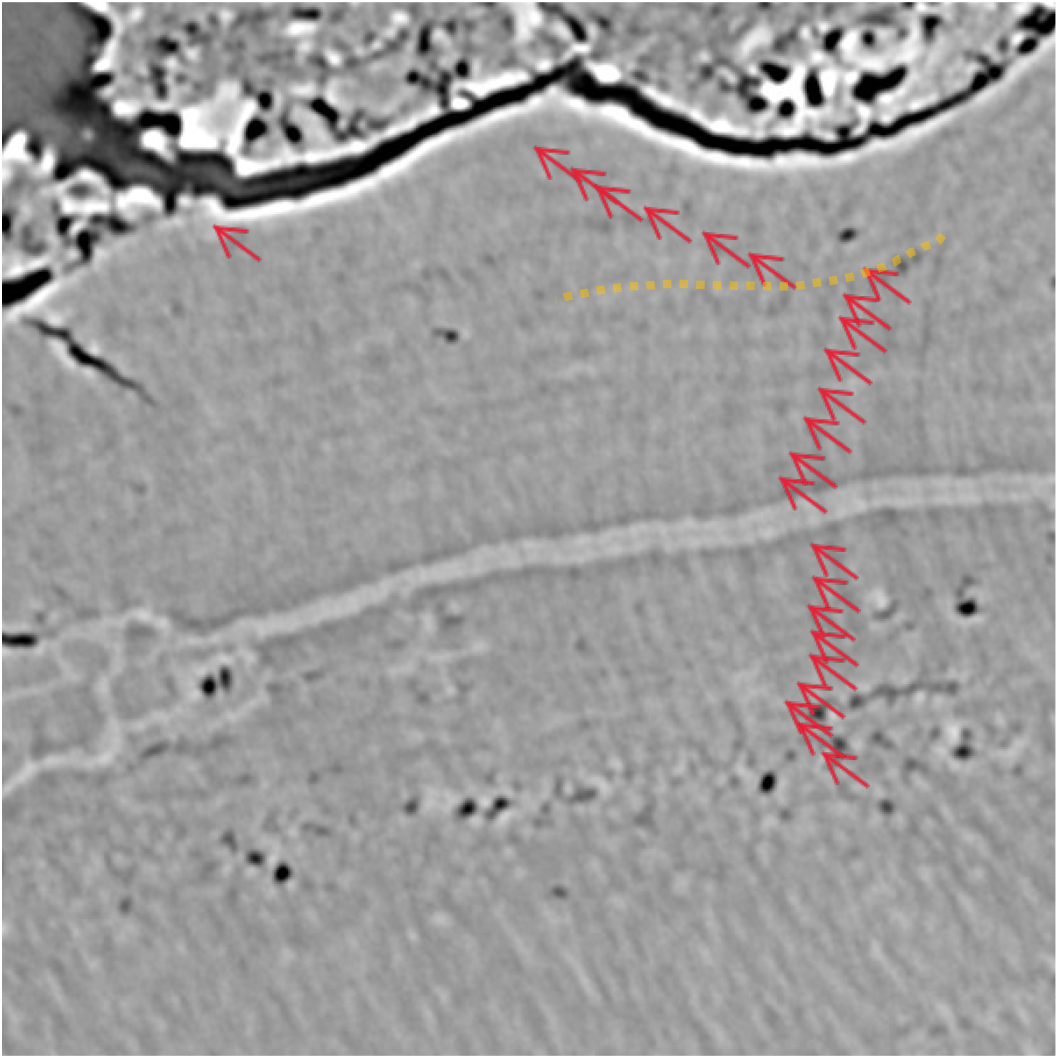
Detailed scan at 6 mm from the root of P^4^. The red arrows indicate the 24 yearly lines. The orange dotted line follows the curvature of the cement lines.

**Figure 4:**
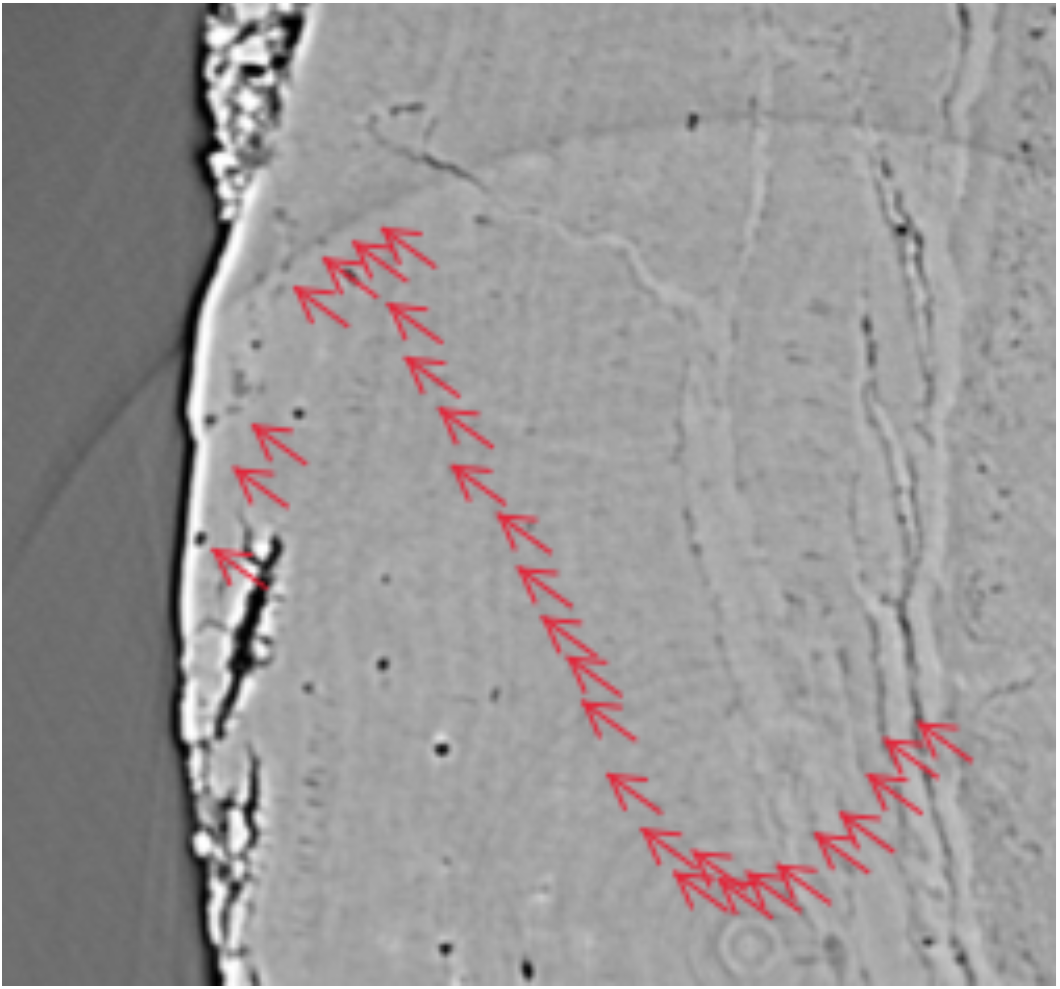
Detailed scan at 0.93 mm from the root of M^1^. The red arrows indicate the 28 yearly lines.

Based on the maps, 10 appropriate positions for the detailed transverse scans were chosen on M^2^ and 4 on P^4^ in areas where acellular cementum was present. Additionally, five suitable positions for the high-resolution transverse scans were chosen on M^1^. All detailed transverse scans on the three teeth were scanned at 40 keV, 70 mm propagation distance, with 3× 0.4 s accumulated images and 3000 projections over 360° (roughly 1 hour per tomogram). In all cases, 40 keV monochromatic radiation was used, with a flux of around 2E11 photons/s/mm^2^. Although acellular cementum is usually utilised for cementochronology (17), the cellular cementum was better preserved in M^1^, and has also been shown to consist of yearly lines (18, 19).

After the detailed scans were completed, we chose the clearest images from the dataset and counted the growth lines manually, since automated counts have been found to underestimate the total number of increments (20). Growth lines were counted on multiple images to assure the maximum number of lines was counted. The lines of Salter were counted multiple times (21) in 2015 and in 2022. In both case the same numbers of lines were found. Representative images are shown here.

## Results

The cement on the root of M^1^ CCH6-c was best preserved out of all three dental elements. The right upper P^4^ CCH6-d is relatively well preserved and displays 24 lines (Figure 3), equivalent to 24 years. The right upper M^1^ displays 28 lines in the tooth root cement (Figure 4), which are equivalent to 28 years. Despite taking detailed scans at 10 positions in M^2^, surface erosion caused the counts to be incomplete at each position, although it was possible to count the yearly lines. The difference between the number of lines of P^4^ and M^1^ is four (=28-24) in the holotype of Homo luzonensis CCH6.

## Discussion

### Development and Age at Death

To assess age at death, the number of lines of Salter need to be added to the age at eruption or the age of root completion. The margins of error of these two approaches are not significantly different, because the timing of these event is equal on a yearly timescale (13).

In *Homo luzonensis* CCH6, the cement of the root of M1 was very well preserved and the cement lines count can be considered reliable. The cement of P4 was slightly less well preserved, but was mostly deformed rather than eroded, since the cement lines follow the shape of the outer surface and the cracks. (Figure 3, orange dotted line).

In anatomically modern humans (*Homo sapiens*), a difference of five lines would be expected between these dental elements, since the M^1^ erupts at age six and the P^4^ at age 11 on average (22). Nevertheless, there is some intraspecific variation and a difference of four lines may be expected in approximately 5% of the population (23). In chimpanzees (*Pan troglodytes*) on the other hand M1 erupts at age three and P^4^ at age seven, so we would expect a difference of four lines between the M^1^ and the P^4^ (22). The mean interval in gorillas (*Gorilla gorilla*) on the other hand would be three lines, since M^1^ erupts in the fourth year of life and P^4^ in the seventh (22). Although individual variation needs to be taken into account (24), cementochronology is the most accurate method for age-at-death estimation (25), and it is possible to speculate that the dental development pattern of *H. luzonensis* might have been more similar to that of extant chimpanzees than to that of anatomically modern humans, although this is not necessarily a reflection of overall development. Even if an unknown number of lines would be missing from P4, this would imply a developmental strategy quite dissimilar from modern humans. Consequently, a reasonable estimate for the eruption ages of M^1^, M^2^ and P^4^ *H. luzonensis* would be three, six and seven years old respectively, based on a chimpanzee-like dental development pattern.

Assuming a chimpanzee-like developmental pattern based on the M^1^-P^4^ eruption interval, the root of M^1^ indicates that CCH6 died at the age of 31, calculated from 28 years of root cement formation and three years of postnatal life before the eruption of M^1^. P^4^, also points to an age at death of 31 years, consisting of 24 years of root cement formation and seven years of postnatal life before the eruption of P^4^. A maximum potential age at death for the type specimen of *H. luzonensis* may be calculated by assuming a human-like developmental pattern with estimated eruption ages of M^1^, M^2^ and P^4^ of six, 12 and 11 years old. This would result in an age at death of 34 or 35 years old. Since the ages calculated assuming a chimpanzee-like developmental pattern correspond perfectly with each other, we think it is likely that the type specimen of *H. luzonensis* was approximately 31 years old at time of death but might have been a few years older.

### Implications for the Life History of *H. luzonensis*

In primates, dental eruption pattern is correlated with brain size (26). The diminutive size of *H. luzonensis* has already hypothesised (8), so, similar to *H. floresiensis*(27), its brain might also have been small, which would not be discordant with the dental eruption pattern found in this study.

In primates, various life history traits, such as life expectancy, are also correlated to the dental eruption pattern (28, 29). In anatomically modern humans, life expectancy has changed over the course of (pre-)history (30). In the Mesolithic, life expectancy was between 21 and 40 years at birth, whereas in the Neolithic life expectancy was 17 to 25 years (31). In the Copper, Bronze and Iron ages, life expectancy at birth was between 28 and 44 years (30). CCH6 died in his or her early thirties. This seems relatively young for current standards but is in line with Mesolithic modern humans, as well as corresponding to the potentially fast development of *H. luzonensis*.

### Possible Evolutionary Implications

*Homo luzonensis*, a small hominin from the Philippines, displays a dental eruption pattern which could be similar to that of modern chimpanzees, but also occurs in a small portion of the modern human population (23). Consequently, its life history traits might to follow a more ape-like pattern than a modern human pattern. This could have implications for human evolution, but more evidence is clearly needed. It had already been established with *H. floresiensis* that relatively recent hominins could have brain sizes in the range of chimpanzees (27).

### Perspectives

Here, we used a new methodology to calculate age at death using synchrotron radiation. This method is non-destructive and can be used on valuable fossils, such as those of Callao Man.The methodology described herein is a proof of concept and holds great potential for the study of fossil hominins and other valuable dental remains that are not suitable for destructive sampling. Although it is difficult to supply conclusive evidence of life history parameters based on two teeth, the methodology could be applied to more complete tooth rows with the prospect of deciphering an individual’s complete dental development. Comparing fossil taxa through time will provide direct evidence for the evolution of dental development with implications for the associated life history traits. Furthermore, this method allows for an assessment of age at death and, when applied to multiple individuals of the same population, might provide an estimate of average life expectancy. As suggested by von Jackowski et al. (15), studying the yearly cementum lines over the whole tooth root cross-section might allow for the detection of potential interruptions or irregularities of the individual layers. These features may be associated with childbirth in females or certain diseases (32–34) and could provide valuable information on the life of fossil hominins. As shown in this work, the potential of using fast, low-resolution measurements combined with slower, high-resolution measurements in selected areas could be important here. To apply cementochronology to the lines of Salter, a standardised protocol for each species, or even each dental element, would be ideal (13). In fossils, however, this would be problematic. Firstly, it would be extremely difficult, if not impossible, to ground-truth any such protocol, due to a lack of fossils with known individual age at death and/or known age at eruption of the respective dental elements. Secondly, due to taphonomical processes, the researcher will have to work with those parts of the tooth cementum that are sufficiently well-preserved. Theoretically, it should be possible to determine whether an individual died in fall/winter or spring/summer. These seasons are characterised by dark or light lines respectively in light microscopy (13). More research on modern humans is needed, however, to determine how the cement layers are formed in a monsoonal climate. It will also be challenging to extrapolate this analysis to synchrotron radiation tomography. It is not currently known how the dark and light lines correspond to the dark and light lines in light microscopy, since the light and dark lines in tomography are caused by differences in density, whereas the cause of the light and dark lines in normal microscopy is still under debate (13) and differs according to the type of microscope used (35, 36).

Furthermore, the strong contrast between cementum and air at the surface generates an intense phase contrast fringe, which interferes with interpretation of near surface lines. This could be partially improved by surrounding the tooth with a medium closer to the electron density of cementum, as in Lak et al. (37).

## Acknowledgements

We thank the National Museum of the Philippines and its Directorial staff for allowing us to scan the specimens and the Cagayan Provincial Government and the Protected Area Management Board-Peñablanca for authorizing fieldwork at Callao Cave. Funding was provided by Muséum national d’Histoire naturelle, the Wenner-Gren Foundation, the Leakey Foundation Research Grant and the National Geographic Society, beamtime was granted by Synchrotron Soleil, and AHvH was supported by a postdoctoral fellowship of the Humboldt Foundation. The micro-CT scans for illustrative purposes were provided by AST-RX, plateforme d’Accès Scientifique à la Tomographie à Rayons X, UAR 2700 2AD Acquisition et Analyse de Données pour l’histoire naturelle CNRS-MNHN, Paris.

